# Whole-Proteome ESM-2 Embeddings Recover Taxonomy and Enable Geometry-Aware Triage of Foodborne Bacterial Genomes

**DOI:** 10.64898/2026.04.26.720952

**Authors:** Jay Gutierrez, Javier Correa Alvarez

## Abstract

Whole-genome sequencing (WGS) has transformed foodborne pathogen surveillance, yet time-sensitive decision-making remains constrained by computationally expensive alignment-centric workflows that scale poorly to outbreak volumes and lack built-in confidence signals. Using 21,657 GenomeTrakr-derived assemblies spanning nine food safety–relevant taxa, we represent each genome by mean-pooling per-protein embeddings from ESM-2 (480 dimensions). The resulting embedding space is dominated by taxonomic structure, exhibiting near-perfect neighborhood consistency for both species and a coarse species/pathotype-derived pathogenicity prior (mean homophily >0.99). Density-based clustering recovered species-coherent structure with high purity and bootstrap stability, while external agreement with the binary pathogenicity prior was only moderate, which is consistent with phylogenetic entanglement by design rather than embedding failure. As a within-genus stress test, kNN separates *E. coli* O157:H7 from non-pathogenic *E. coli* with ∼98% accuracy (5-fold CV), demonstrating that known pathotype annotations are preserved in the embedding geometry even among closely related genomes. We position this mean-pooling baseline relative to contextual genome language models that retain protein order or operon-scale context, and outline how embedding geometry (homophily, purity, outliers) can serve as a principled confidence layer in bio-surveillance-oriented triage pipelines.

## 1. Introduction

Foodborne diseases remain a major global burden, with World Health Organization estimates of ∼600 million illnesses and ∼420,000 deaths per year attributable to unsafe food [1]. Modern bio-surveillance increasingly relies on whole genome sequencing (WGS), and networks such as FDA’s GenomeTrakr have enabled more precise traceback and outbreak investigation at scale [2]. Despite this progress, classical approaches to bacterial genome comparison span a spectrum of resolution and computational cost. At the coarsest level, whole-genome similarity scores (such as average nucleotide identity) provide a single pairwise measure suitable for species-level classification but insensitive to strain-level variation relevant to outbreak tracking. Gene-based typing schemes offer standardized, portable strain identifiers by comparing sets of conserved genes across isolates, but these depend on curated reference databases that require ongoing maintenance and may lag behind novel diversity.

Alignment-free methods based on short sequence patterns, including MinHash sketches as implemented in Mash [3], dramatically reduce computational cost but primarily capture nucleotide-level similarity without explicitly encoding functional or structural signals relevant to pathogenicity. High-resolution variant-calling pipelines provide the finest discrimination for outbreak cluster definition but require reference genome selection, read mapping, and base-by-base comparison, a workflow poorly suited to real-time triage of novel or divergent isolates. In this context, protein language model embeddings offer a complementary representation: alignment-free by construction, implicitly encoding evolutionary and functional signals learned from massive sequence databases, and amenable to geometric analyses that can surface confidence and novelty signals absent from discrete typing schemes [4,5].

In parallel, protein language models (PLMs) such as ESM have shown that structure and function can emerge from self-supervised learning on large protein sequence corpora [4,5]. This has motivated a wave of genome-scale modeling efforts that treat genomic content as language [6], either at the nucleotide level (DNABERT [10], DNABERT-2 [11], Nucleotide Transformer [9], HyenaDNA [12], GenSLMs [13]) or at the protein/gene level (gLM [7], Protein Set Transformer [8], BacPT [18], Bacformer [19]).

In this landscape, a practical question remains: how much can we gain from the simplest possible genome representation built from PLMs, i.e., mean pooling per-protein embeddings, when scaled to tens of thousands of genomes under realistic bio-surveillance constraints? And can the geometry of the embedding space be turned into actionable signals for operational systems (confidence, novelty, drift, anomaly triage), rather than treated as mere visualization artifacts?

## Contributions

This manuscript provides a large-scale, geometry-first characterization of whole-proteome ESM-2 embeddings for foodborne surveillance. Specifically, we (i) implement a cache-first embedding pipeline that enables reproducible reuse across analyses and downstream models; (ii) quantify the embedding space’s intrinsic dimensionality and clustering geometry; (iii) introduce homophily, purity, and outlier scores as operationally relevant confidence signals; and (iv) position this mean-pooling baseline in a transparent way relative to contextual genome language models designed to model operon- and genome-scale dependencies.

## 2. Materials and Methods

### 2.1 Dataset and Labeling

We analyze 21,657 bacterial genomes drawn from a GenomeTrakr-derived collection (https://www.ncbi.nlm.nih.gov/pathogens/) and grouped into nine taxa [2]:

| Taxon | n | Label |
| --- | --- | --- |
| *Salmonella enterica* | 7,000 | pathogenic |
| *Listeria monocytogenes* | 4,500 | pathogenic |
| *E. coli* (non-pathogenic group) | 4,312 | non-pathogenic |
| *Bacillus subtilis* | 2,361 | non-pathogenic |
| *E. coli* O157:H7 | 1,500 | pathogenic |
| *Citrobacter koseri* | 897 | non-pathogenic |
| *Listeria innocua* | 449 | non-pathogenic |
| *Escherichia fergusonii* | 438 | non-pathogenic |
| *Citrobacter freundii* | 200 | non-pathogenic |

Because isolate-level virulence phenotypes are not uniformly available, coarse pathogenicity prior was derived from species/pathotype assignments. This label is intentionally treated as a *bio-surveillance prior*, not as ground truth for every individual isolate; it is suitable for studying embedding geometry, retrieval consistency, and coarse triage, but it cannot resolve within-species virulence variation.

Accordingly, within-genus stress tests (e.g., *E. coli* O157:H7 vs non-pathogenic *E. coli*) are reported, and performance is interpreted as pathotype-level discriminability under known annotations, rather than as discovery of novel virulent isolates in the absence of prior labeling.

### 2.2 Whole-Proteome Embeddings (ESM-2 Mean Pooling)

Protein sequences were extracted from GenBank annotations. Each protein was embedded with ESM-2 and the pretrained `esm2_t12_35M_UR50D`[5] and reduced to a single 480-dimensional vector by mean pooling over token embeddings. Genome embeddings were then computed as mean pooling across all proteins in the genomes.

This “bag-of-proteins” genome representation is deliberately simple, and corresponds to the context-free baselines used in operon-/contig-scale genomic language modeling (e.g., mean pooling of ESM-2 protein embeddings prior to contextual modeling in gLM) [7].

### 2.3 Cache-First Execution and Reproducibility

Content-addressed caching of per-genome protein embedding tensors was used, where keys incorporate the model ID, maximum protein length of 1024, pooling configuration, and a hash of input sequences. Cache hits were made deterministic across runs and environments by this design, sharded embedding generation across accelerators was enabled, and downstream analyses and models were decoupled from recomputation.

Large-scale embedding runs were executed on the EAFIT Apolo-3 HPC cluster using 2-way sharding across dual NVIDIA H100 NVL GPUs (one worker per GPU). Subsequent analytics and figure generation were CPU-feasible using the pooled embedding bundle. Moreover, for analysis reproducibility, pooled embeddings and metadata were exported as a compressed NPZ bundle and available in the repository (see below).

### 2.4 Dimensionality Reduction and Clustering

Embeddings are standardized (zero mean, unit variance per dimension). We compute PCA (50 components) and report explained variance. For visualization we compute UMAP (`n_neighbors=30, min_dist=0.1`) [14] and t-SNE (`perplexity=30`) [20] from PCA-reduced features.

For clustering we run HDBSCAN (`min_cluster_size=50, min_samples=10`) on the first 20 PCs [15]. Internal cluster validity is quantified by silhouette score, Calinski–Harabasz index, and Davies–Bouldin index. Cluster purity is defined as the fraction of cluster members sharing the dominant label (either species or pathogenicity prior). External agreement with the binary pathogenicity prior is reported using ARI and NMI. We assess cluster stability via bootstrap resampling and cluster-wise Jaccard matching (50 iterations; 80% subsampling) following Hennig [16].

### 2.5 Homophily as a Geometry-Aware Confidence Signal

We compute kNN homophily (k=20 unless otherwise stated) as the fraction of a genome’s k nearest neighbors (Euclidean distance in standardized embedding space) sharing its label, using both species labels (taxonomy consistency) and the binary pathogenicity prior (coarse risk consistency). Further, we additionally compute multi-scale homophily curves (k ∈ {1, 5, 10, 20, 50, 100}) to characterize how label consistency changes with neighborhood radius.

### 2.6 Outlier Detection

We operationalize outliers using two complementary criteria: distance outliers are genomes whose k-neighbor distance in 20D PCA space exceeds Q3 + 1.5×IQR, while cluster noise are genomes labeled as noise by HDBSCAN (cluster = −1). We report “strict” outliers as genomes satisfying both criteria (high-confidence anomalies).

### 2.7 Within-Genus Stress Tests

To partially mitigate label–taxonomy confounding, we report two focused stress tests in which pathogenic and non-pathogenic priors occur within a related genus-level neighborhood. Concretely, we analyze *Listeria* (*L. monocytogenes* vs *L. innocua*) and *Escherichia* (*E. coli* O157:H7 vs non-pathogenic *E. coli*). For each subset we report kNN vote accuracy (k=20), balanced accuracy, mean homophily, and silhouette score. Together, these methods operationalize the framework depicted in Figure 1 and enable the geometry-aware triage signals we report below.

**Figure 1.**
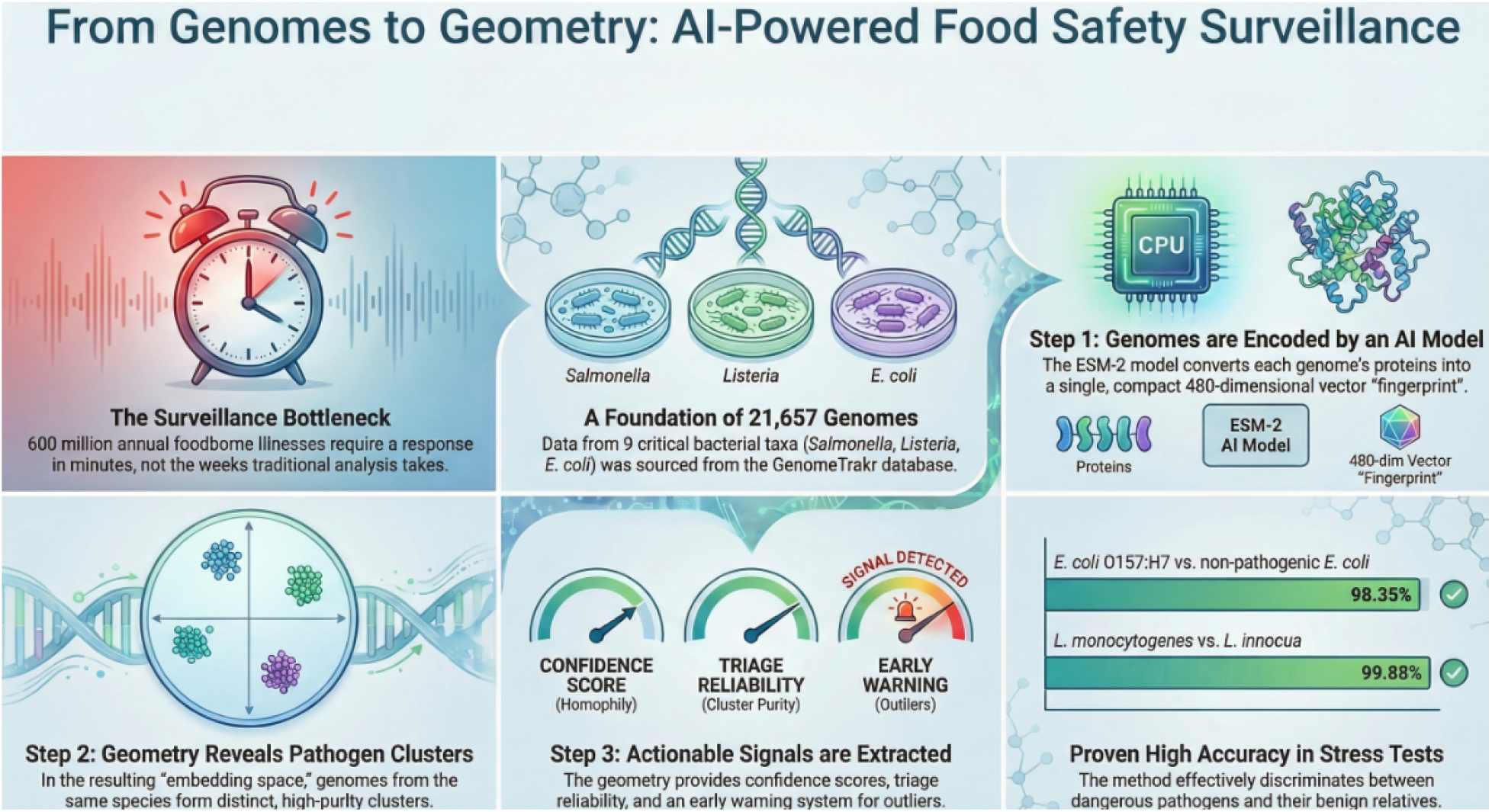
Conceptual overview of the embedding-based surveillance framework. Schematic of the embedding-based triage framework for foodborne bacterial surveillance. The pipeline addresses the surveillance bottleneck where 600 million annual foodborne illnesses demand rapid response that traditional alignment-based workflows cannot deliver. A foundation of 21,657 GenomeTrakr-derived genomes from nine critical taxa (*Salmonella, Listeria, E. coli*, and others) is processed through ESM-2, which converts each genome’s proteins into a compact 480-dimensional vector “fingerprint” (Step 1). In the resulting embedding space, genomes from the same species form distinct, high-purity clusters that reveal taxonomic geometry (Step 2). From this geometry, three actionable signals are extracted (Step 3): confidence scores (homophily), triage reliability (cluster purity), and early warnings (outliers). The framework achieves high accuracy in stress tests discriminating dangerous pathogens from their benign relatives (98.35% for *E. coli* O157:H7 vs non-pathogenic *E. coli*; 99.88% for *L. monocytogenes* vs *L. innocua*), demonstrating operational utility for rapid, alignment-free pathogen triage.

### 2.8 Robustness to Proteome Incompleteness (Protein Dropout Validation)

To evaluate resilience to incomplete assemblies and contamination, we run three cache-only stress tests on a balanced sample of 200 genomes per species (9 taxa; n=1,800). (i) Protein dropout: randomly retain 90%, 75%, or 50% of proteins. (ii) Contig dropout: randomly retain 90%, 75%, or 50% of contigs as a fragmentation proxy. (iii) Contamination mixing: replace 5%, 10%, or 20% of proteins with proteins sampled from a different species (0% as control). Each rate is repeated three times, yielding n=5,400 per rate. For each perturbed embedding we compute cosine similarity to the full-proteome embedding, kNN neighborhood stability (Jaccard overlap; k=20), changes in species/pathogenicity homophily, and outlier rate based on the reference kNN distance threshold (Q3 + 1.5×IQR). This analysis is CPU-feasible on Apolo-3 using the cached `prot_emb_*.pt`files and is fully reproducible via `notebooks/foodguard_simulated_drift_study.ipynb`.

## 3. Results

We present results organized around the three operational signals highlighted in Figure 1: embedding geometry and clustering (Sections 3.1–3.5), confidence via homophily (Section 3.6), and outlier detection for quality control (Section 3.7). Within-genus stress tests (Section 3.8) validate discriminability under challenging conditions.

### 3.1 Low Dimensionality of Mean-Pooled Embeddings

PCA indicates that the embedding space is dominated by a small number of factors: PC1 explains 76.56% of variance, PC2 explains 10.95%, and only 3 PCs are needed to capture 90% of variance (Figure 2A). This strong low-dimensional structure is consistent with broad taxonomic and proteome-level signals shaping the dominant geometry of the space.

**Figure 2.**
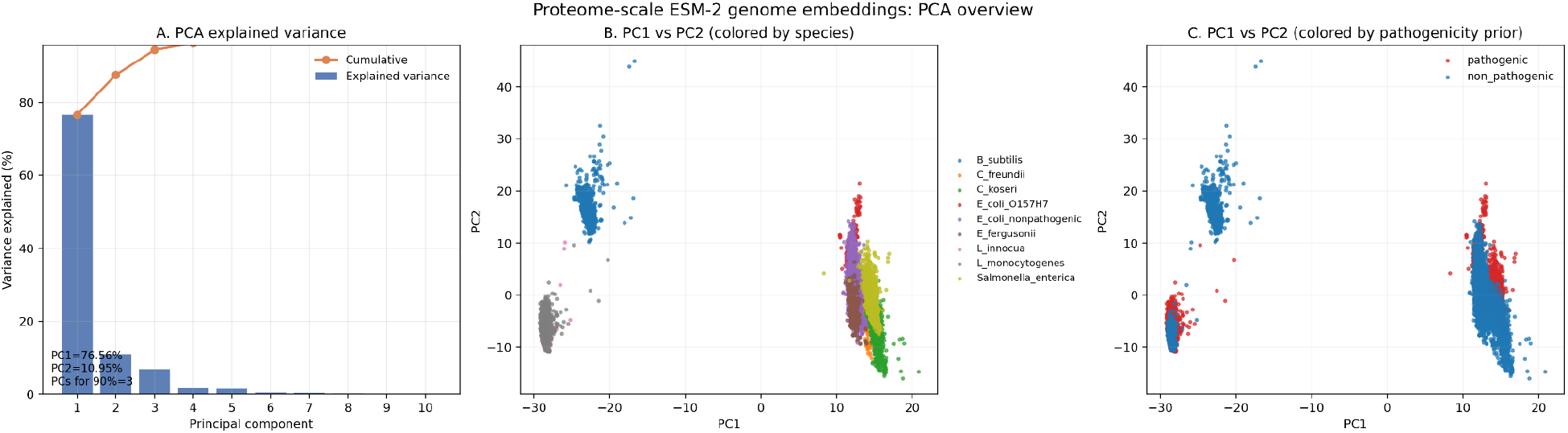
PCA overview of whole-proteome genome embeddings (explained variance; PC1 vs PC2 colored by species and pathogenicity prior). Whole-proteome ESM-2 genome embeddings projected with PCA after standardizing embedding dimensions. (A) Explained variance ratio for the first 10 principal components (bars) with cumulative variance (line); PC1 explains 76.56% of variance, PC2 explains 10.95%, and 3 PCs exceed 90% cumulative variance (94.3%). (B) PC1 vs PC2 for all genomes (n=21,657), colored by species. (C) Same PC1 vs PC2 projection colored by the pathogenicity prior used in this study (species/pathotype-derived). Each point represents one genome embedding obtained by mean-pooling per-protein ESM-2 representations (480 dimensions) computed from GenBank-annotated protein sequences.

Importantly, this result should be interpreted as a descriptive property of the current dataset and representation (standardized, mean-pooled protein embeddings) rather than as a claim that the underlying biology is intrinsically three-dimensional. PCA is a global linear model: it concentrates variance along directions that best explain the dataset under a linear assumption, and additional biologically meaningful structure can reside in higher-order PCs and in non-linear manifolds revealed by methods such as UMAP and t-SNE (Figure 3).

**Figure 3.**
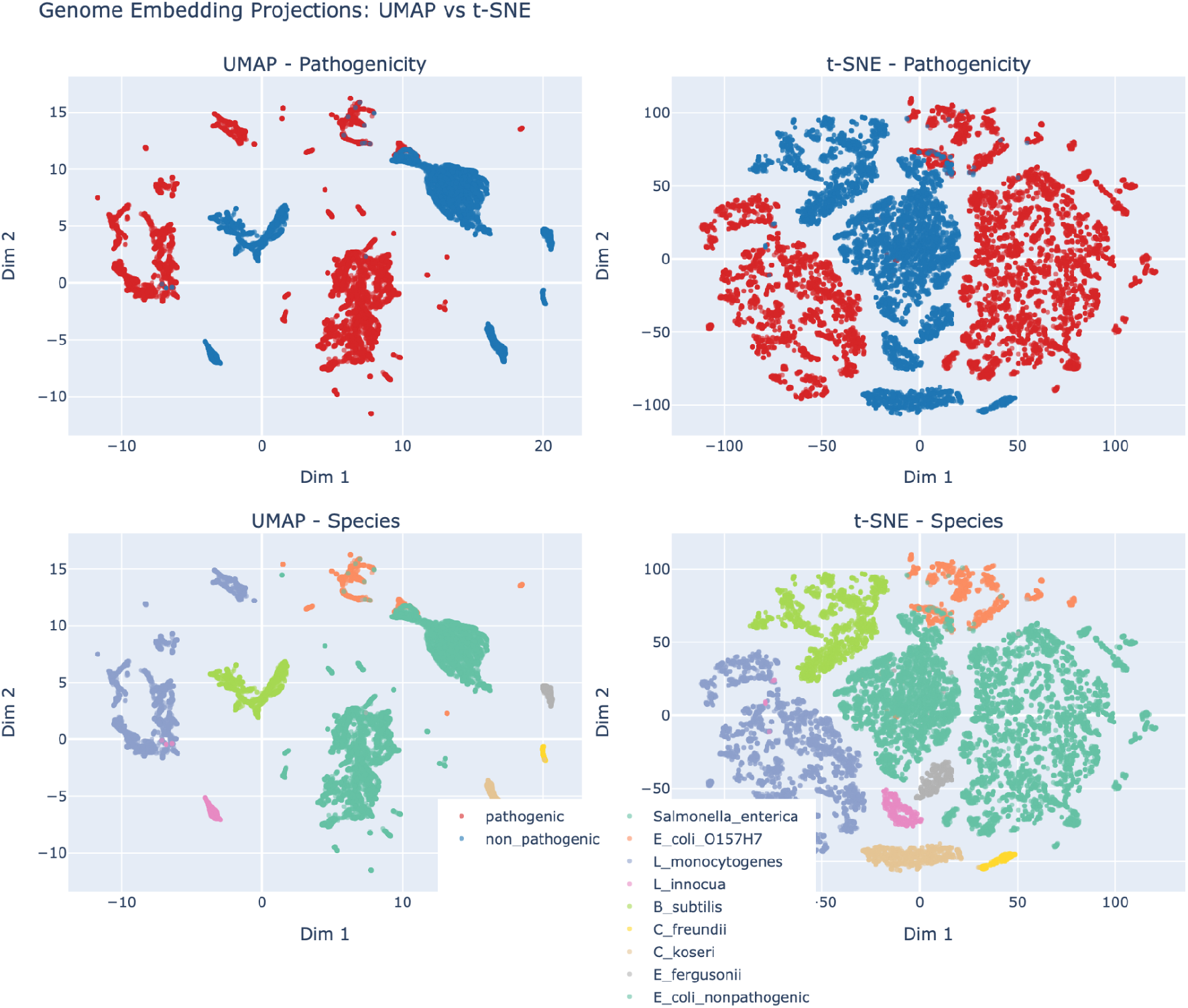
UMAP and t-SNE projections by pathogenicity prior and species. Non-linear 2D projections of whole-proteome genome embeddings, computed after standardization and PCA pre-reduction (50 components). Top row: UMAP [14] (left) and t-SNE [20] (right) colored by the pathogenicity prior (species/pathotype-derived). Bottom row: the same UMAP and t-SNE projections colored by species (nine taxa). Across both methods, genomes form largely species-coherent islands, while the pathogenicity prior aligns with these islands due to phylogenetic confounding; mixed regions highlight closely related taxa with different priors and motivate confidence-aware retrieval and context-aware modeling. Axes are arbitrary, and global distances/layout are not interpreted as metric-preserving.

### 3.2 Taxonomic Structure in Low-Dimensional Projections

PCA already reveals pronounced taxonomic structure in the first two axes (Figure 2B–C), and non-linear projections via UMAP and t-SNE sharpen the same species-level islands (Figure 3).

Qualitatively, Figure 2B suggests that the dominant PCA axis separates broad phylogenetic regimes: genomes from Firmicutes in this dataset (e.g., *Bacillus* and *Listeria*) occupy the negative PC1 region, while Enterobacterales (e.g., *Salmonella, Escherichia, Citrobacter*) concentrate at positive PC1 values. Within the Enterobacterales region, the spread along PC2 and the partial overlap between closely related *Escherichia* taxa visually anticipate the downstream pattern we quantify with homophily and within-genus stress tests: most neighborhoods are internally consistent, but a small subset of genomes sits near decision boundaries where retrieval-based triage becomes less reliable.

Figure 3 reinforces this interpretation. The species-colored views confirm that the “island” structure is robust across both UMAP and t-SNE, with the same local groupings emerging under different projections, indicating that the observed clusters are not artifacts of PCA alone. Meanwhile, the pathogenicity-colored views (Figure 2C and Figure 3, top row) illustrate why we treat the binary label as a prior rather than isolate-level ground truth: the red/blue separation largely tracks taxonomy (because the label is species/pathotype-derived), and the most informative regions are the mixed neighborhoods where related taxa carry different priors (e.g., *Escherichia* groups and the *Listeria* pair).

We emphasize that UMAP and especially t-SNE are visualization tools that prioritize local neighborhood structure and can distort global geometry. Accordingly, we interpret Figure 3 in terms of neighborhood consistency and cross-method stability rather than in terms of absolute inter-island distances or global layout.

### 3.3 Density-Based Clustering and Stability

Consistent with these projections, HDBSCAN identifies 34 clusters with 35.6% noise points, a typical outcome for density-based clustering in biological embedding spaces with variable density. Internal validation indicates strong cluster structure (excluding noise points): silhouette 0.555, Calinski–Harabasz 10,849.94, and Davies–Bouldin 1.0719. As a sanity check, silhouette drops well below zero under permutation of cluster labels (−0.229; 5,000-point sample; excluding noise), confirming that the observed value reflects non-trivial structure rather than a clustering artifact.

Bootstrap resampling with cluster-wise Jaccard matching (50 iterations; 80% subsampling) indicates that most clusters are robust to resampling, with mean Jaccard stability 0.81 overall and 23/34 clusters classified as stable or highly stable (mean Jaccard ≥ 0.75).

### 3.4 Species Centroid Distances as Confusability Priors

At a coarser resolution, centroid-to-centroid distances between species provide an interpretable map of which taxa are closest in this embedding space and therefore most likely to generate boundary cases under retrieval-based triage (Figure 4). The matrix recapitulates the dominant taxonomic geometry: *Bacillus* and *Listeria* centroids are far from the Enterobacterales taxa (typical distances ∼40–46), consistent with the broad PC1 separation in Figure 2B. Within each regime, distances compress and identify the most confusable neighborhoods. The closest pair is the *Listeria* duo (*L. monocytogenes* vs *L. innocua*; 4.04). Within *Escherichia*, the non-pathogenic *E. coli* centroid lies near *E. fergusonii* (5.34) and the O157:H7 pathotype remains relatively close to both (e.g., *E. coli* O157:H7 vs non-pathogenic *E. coli*; 12.95). The two *Citrobacter* species are also closer to each other than to most other taxa (10.46). Notably, some cross-genus Enterobacterales distances remain small compared to the Gram-positive/Gram-negative gap (e.g., *Salmonella enterica* is 8.02 from *E. fergusonii* and 8.31 from non-pathogenic *E. coli*), implying that species identity is the primary organizing signal while fine-grained separability is uneven across taxa.

**Figure 4.**
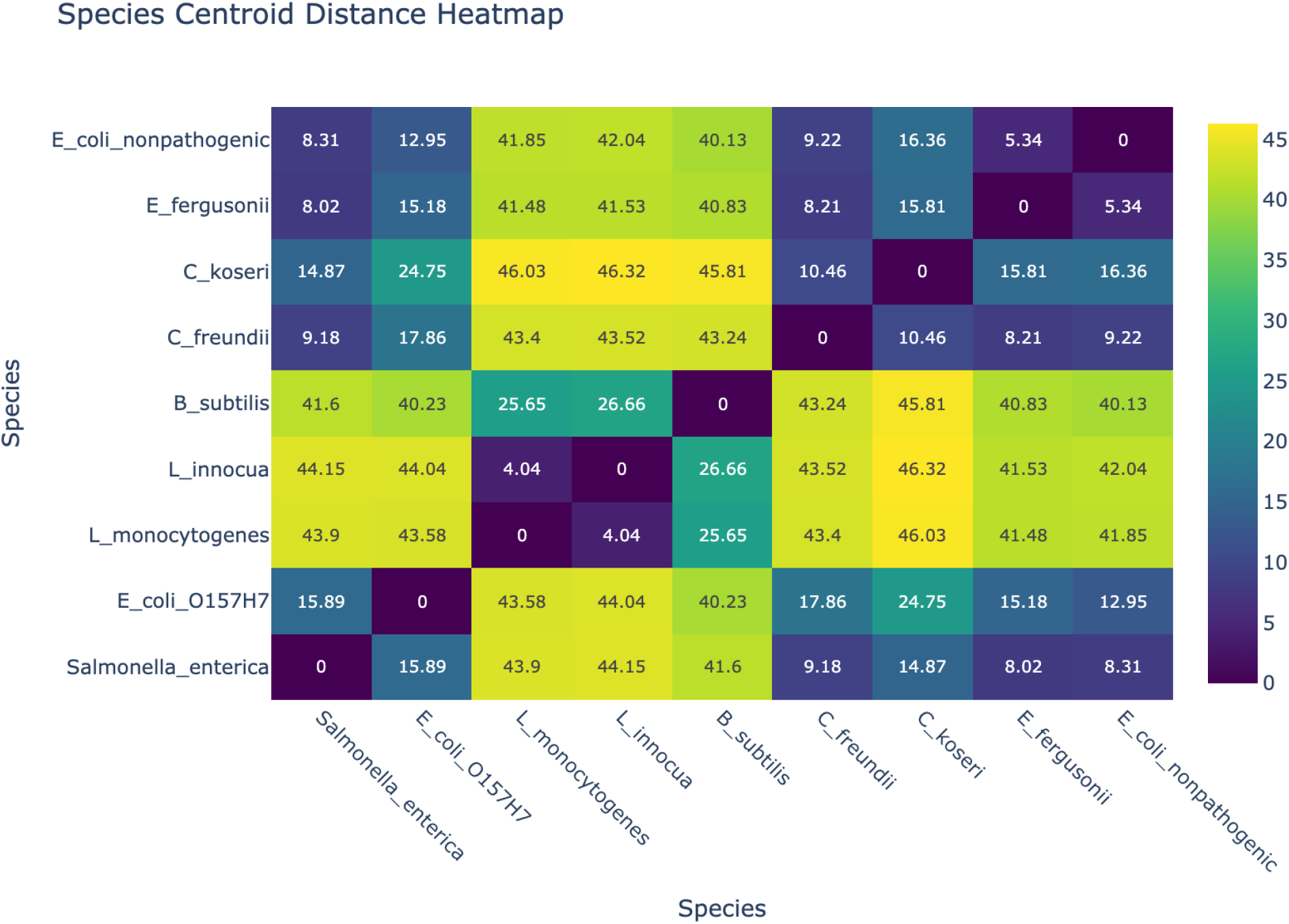
Species centroid distance heatmap (Euclidean distance in standardized embedding space). Pairwise Euclidean distances between species centroids in the standardized whole-proteome embedding space. Each species centroid is the mean of its z-scored genome embeddings (mean-pooled 480-dimensional ESM-2 representations), and each cell reports the L2 distance between two species centroids (diagonal = 0). Smaller distances indicate taxa whose *average* embeddings occupy nearby regions of the space and are therefore more likely to yield near-boundary neighbors under kNN retrieval; larger distances reflect broad taxonomic separation. Because centroids compress within-species heterogeneity, the heatmap is interpreted as a coarse summary of global geometry rather than as a distribution-aware or phylogenetically calibrated distance.

These centroid distances should be read as a pragmatic “confusability map” rather than as a calibrated evolutionary distance. Averaging collapses within-species multimodality and is sensitive to unequal sampling and within-species variance, and Euclidean distances in standardized embedding space depend on the chosen scaling and metric. Accordingly, we use Figure 4 to motivate which within-genus or within-order comparisons deserve conservative handling downstream, not to claim a universal notion of biological distance.

To validate centroid distances as confusability priors, we quantified cross-species neighbor mixing (k=20) and found a strong inverse association between centroid distance and symmetric mixing (Spearman ρ = −0.65; p = 1.5×10^−5^; Figure 7). Moreover, bootstrap confidence intervals for key close pairs are narrow (e.g., *Listeria* centroid distance 4.04, 95% CI 3.99–4.12; O157:H7 vs non-pathogenic *E. coli* 12.95, 95% CI 12.71–13.21), supporting that these confusability relationships are stable given current sampling.

### 3.5 Pathogenicity Separation and Phylogenetic Entanglement

Binary pathogenicity priors show substantial centroid separation (multivariate effect size 7.52; distance ratio inter-/intra-class 1.106), but external agreement between clusters and pathogenicity is only moderate (ARI 0.2217; NMI 0.3429). At first glance, this may appear inconsistent with high cluster purity, yet the metrics answer different questions. High purity indicates that most clusters are dominated by a single prior label, whereas ARI/NMI penalize over-partitioning: splitting one label into many label-pure clusters (e.g., multiple *Salmonella* subclusters) yields low global agreement with a two-class scheme even when each subcluster is internally homogeneous. In short, high purity with low ARI/NMI reveals that taxonomy, and within-taxonomy substructure, is the primary organizing signal, and that pathogenicity is entangled with phylogeny by design rather than cleanly separable as a binary attribute.

Despite this confounding, cluster membership is still actionable for triage: 33/34 clusters (97%) achieve ≥0.90 pathogenicity purity (and similarly ≥0.90 dominant-species purity), supporting a cluster-based confidence layer. We further stress-tested generalization by withholding each taxon and predicting its pathogenicity prior from the remaining taxa using kNN. As expected under a species/pathotype-derived prior, zero-shot transfer across taxa is unreliable (several taxa flip labels when held out), reinforcing that our binary labels are useful for retrieval consistency and coarse routing within a known reference set, not for “unknown taxon” pathogenicity inference.

### 3.6 Homophily and Boundary Cases

Mean homophily is near-perfect for both labels (k=20): pathogenicity homophily 0.9929 and species homophily 0.9923.

Boundary cases remain informative: 102 genomes have pathogenicity homophily < 0.5, and 111 genomes have species homophily < 0.5. These low-homophily cases concentrate primarily in *Escherichia* groups (O157:H7 and non-pathogenic *E. coli*), rather than in taxa that are globally well-separated (e.g., *Salmonella*).

Multi-scale homophily curves (Figure 5) show that neighborhood agreement remains high across scales for the dataset as a whole (mean homophily 0.9958 at k=1 and 0.9862 at k=100), but the degradation is concentrated in confusable taxa. In particular, *E. coli* O157:H7 shows a larger drop (0.973 at k=1 to 0.934 at k=100) and a substantial rise in mixed neighborhoods as k grows: at k=100, 20.1% of O157:H7 genomes have homophily < 0.9. Conversely, taxa that are geometrically isolated in the global embedding space (e.g., *Salmonella enterica* and *L. monocytogenes*) remain near-perfect even at large k. Interpreted conservatively, Figure 5 operationalizes “hardness” as a scale-dependent property of local geometry, and motivates using homophily-derived confidence to identify when retrieval-based triage should defer to richer, context-aware modeling.

**Figure 5.**
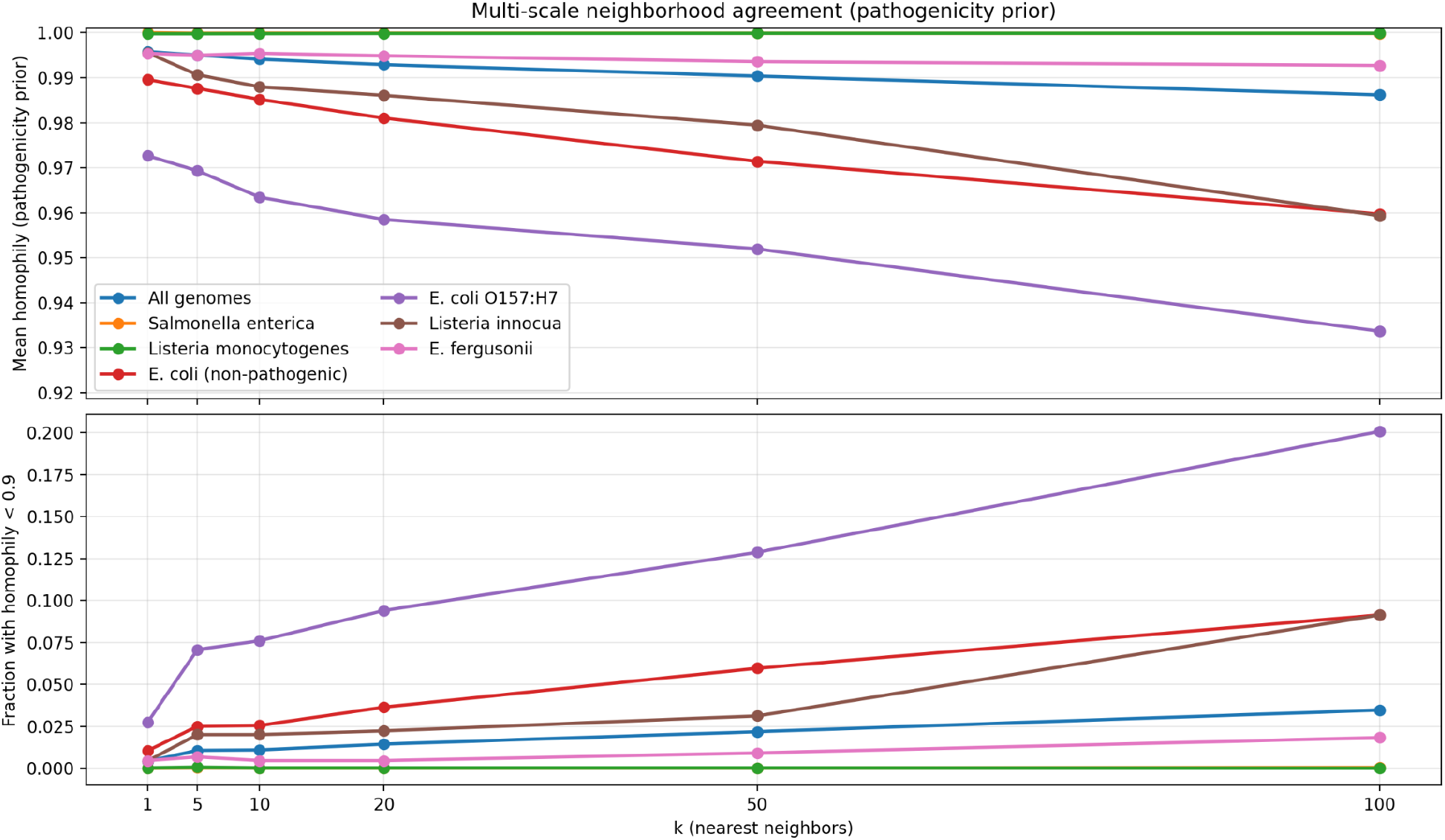
Multi-scale homophily of the pathogenicity prior (mean agreement and mixed-neighborhood fraction vs k). Multi-scale neighborhood agreement for the pathogenicity prior (species/pathotype-derived). For each genome, pathogenicity homophily is defined as the fraction of its k nearest neighbors (Euclidean distance in the standardized 480-dimensional embedding space) that share the same prior label. Top: mean homophily vs k for all genomes and selected taxa; the y-axis is intentionally zoomed to reveal sub-ceiling differences. Bottom: fraction of genomes with mixed neighborhoods, operationally defined here as homophily < 0.9 (a high threshold chosen to highlight early mixing), as k increases. Because the prior is partially confounded with taxonomy, high homophily primarily reflects taxonomic coherence of the embedding space; the informative signal is the subset of genomes whose local neighborhoods become label-mixed as the neighborhood radius expands, highlighting boundary cases where confidence-aware triage is warranted.

### 3.7 Outliers as QC and Novelty Candidates

Outlier analysis yields 510 distance outliers (2.4%) and 7,707 HDBSCAN noise points (35.6%); their intersection defines 252 strict outliers (1.2%). Strict outliers are enriched in *E. coli* (both groups), *E. fergusonii*, and *Salmonella*, with extreme k-neighbor distances (up to 116.39 in PCA space). These cases are suitable for targeted QC review (assembly completeness, contamination, annotation artifacts) or as candidates for novelty/drift monitoring in operational bio-surveillance settings.

Using available assembly statistics as a lightweight QC proxy, strict outliers tend toward larger genomes and proteomes (median genome size 5.18 Mb vs 4.78 Mb; median proteins 4,902 vs 4,403) and exhibit a heavy-tailed fragmentation profile (75th percentile contig count 231 vs 54). This heterogeneity is consistent with a mixture of biologically larger genomes, annotation variation, and genuinely problematic assemblies, thus reinforcing that outlier flags should trigger review rather than be treated as definitive novelty calls.

### 3.8 Within-Genus Stress Tests and Confidence Calibration

Although the full dataset is dominated by taxonomy, within-genus subsets still show meaningful separation. We emphasize that these experiments measure discrimination under known genus/pathotype annotations (e.g., O157:H7 vs non-pathogenic *E. coli*), not discovery of novel virulent isolates in the absence of prior labels.

| Subset | n | kNN(20)<br>accuracy<br>(5-fold CV) | Balanced<br>accuracy<br>(5-fold CV) | Mean<br>homophily | Silhouette |
| --- | --- | --- | --- | --- | --- |
| *Listeria* (*L. monocytogenes* vs *L. innocua*) | 4,949 | 0.9992 | 0.9955 | 0.9986 | 0.180 |
| *E. coli* (O157:H7 vs non-pathogenic) | 5,812 | 0.9842 | 0.9830 | 0.9752 | 0.392 |

For the *E. coli* stress test, 5-fold CV accuracy is 0.984 (95% CI 0.981–0.987) with balanced accuracy 0.983 (95% CI 0.979–0.987), confirming that performance is not an artifact of a single split. This indicates that even the mean-pooled baseline can support rapid, retrieval-based triage among related taxa, while also revealing where confusions persist (particularly within *Escherichia*). Moreover, beyond point accuracy, Figure 6 provides an actionable calibration view for bio-surveillance-oriented triage: in 5-fold CV on the *E. coli* subset, deferring the lowest-consensus neighborhoods (kNN vote fraction threshold 0.9; 4.7% deferral) captures 71.7% of errors and improves covered-set accuracy to 99.53%, while a stricter threshold (0.98; 8.9% deferral) captures 87.0% of errors and yields 99.77% accuracy on the non-deferred set. This illustrates how neighborhood agreement can be converted into an operator-facing “defer vs decide” policy with measurable trade-offs.

**Figure 6.**
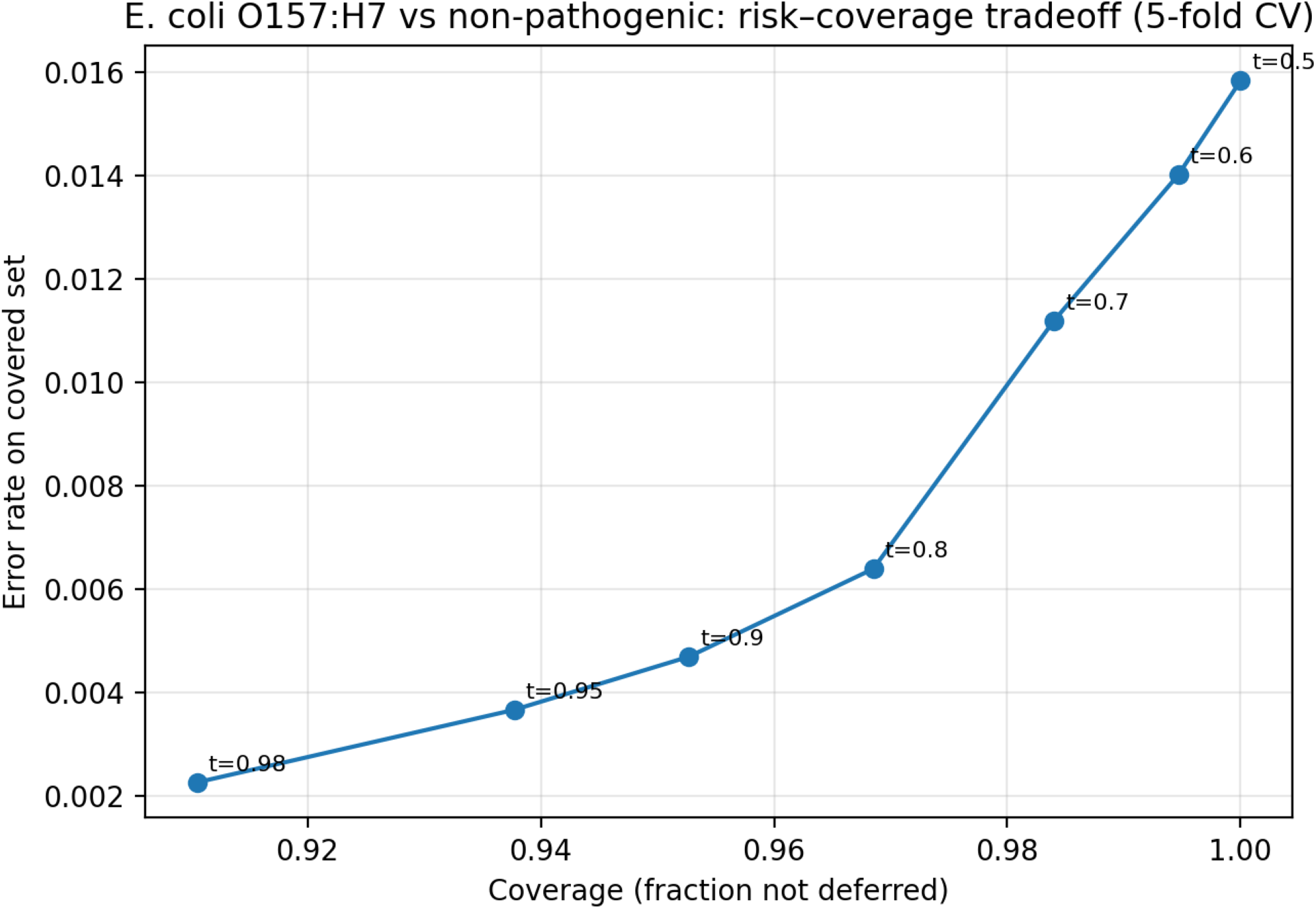
Within-genus confidence calibration (risk–coverage tradeoff) for *E. coli* O157:H7 vs non-pathogenic *E. coli*. Risk–coverage tradeoff for within-genus kNN classification of *E. coli* O157:H7 vs non-pathogenic *E. coli* in the standardized 480-dimensional embedding space (k=20), evaluated with 5-fold stratified cross-validation. Each point corresponds to a vote-fraction threshold on the predicted class; genomes below the threshold are deferred. Coverage is the fraction not deferred, and risk is the error rate on the non-deferred set. This converts neighborhood agreement into an operator-facing “defer vs decide” policy with measurable trade-offs.

**Figure 7.**
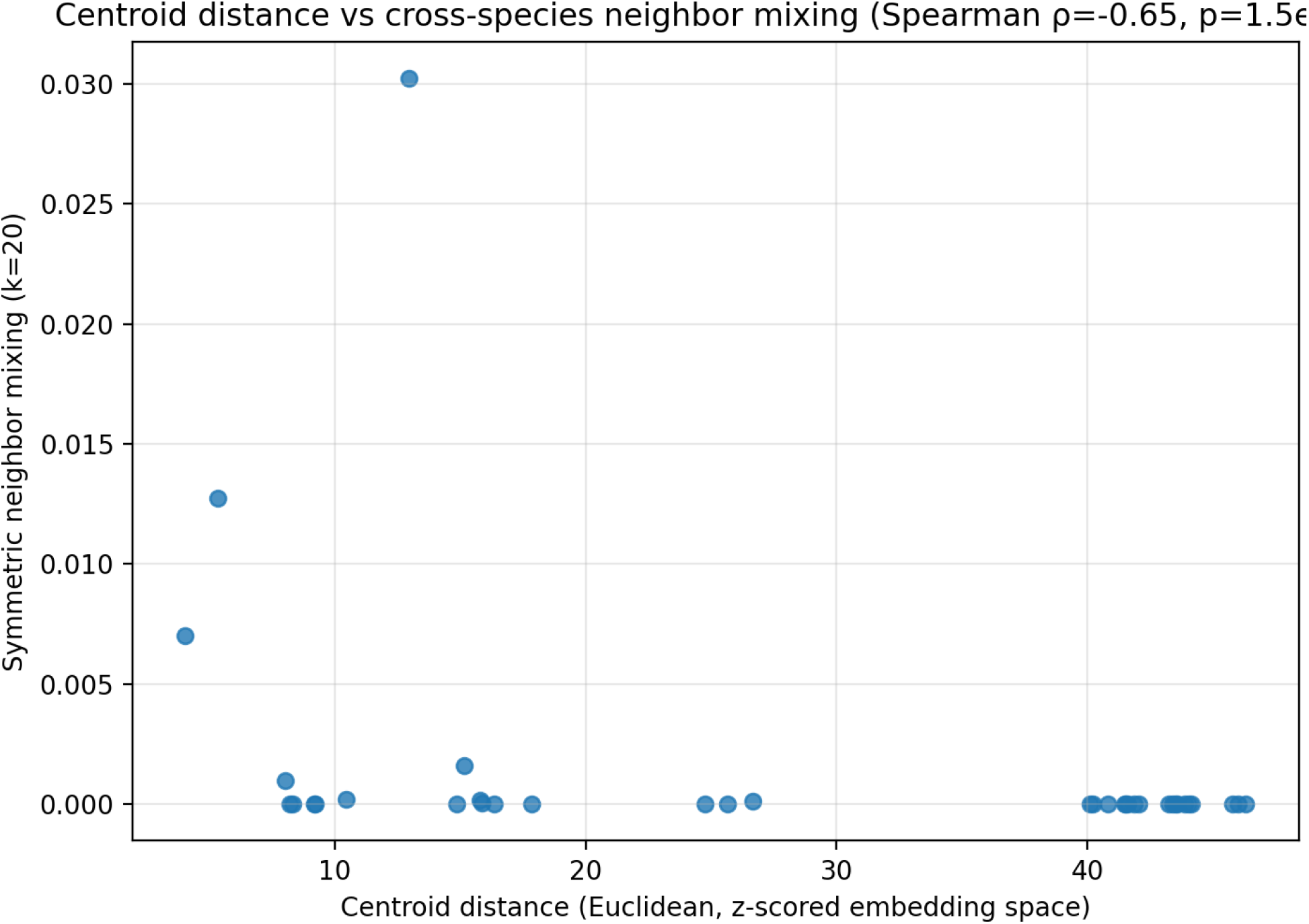
Validating centroid distances as confusability priors. Relationship between inter-species centroid distance (Euclidean in standardized embedding space) and symmetric cross-species neighbor mixing (k=20), computed across all species pairs. Each point represents one taxon pair. Lower centroid distances correspond to higher mixing, supporting centroid distance as a coarse confusability prior (Spearman ρ = −0.65; p = 1.5×10^−5^). Mixing is symmetrized to reduce the effect of sampling imbalance on directional neighbor rates.

### 3.9 Robustness to Proteome Incompleteness

Robustness validation under missingness and contamination shows a consistent pattern: global embeddings remain highly stable while local neighborhoods degrade, and outlier rates rise sharply under severe perturbation (Figure 8). Under protein dropout, cosine similarity remains near 1.0 even at 50% retention (median 0.999981; IQR 0.999975–0.999985), but kNN Jaccard drops from 0.739 (IQR 0.667–0.905) at 90% retention to 0.667 (0.481–0.739) at 75% and 0.481 (0.333–0.600) at 50%. Outlier rate increases from a 2.6% baseline to 3.1% (90%), 4.6% (75%), and 26.4% (50%). Contig dropout is more disruptive: at 90% retention, Jaccard is 0.818 (0.481–1.000) with 7.8% outliers, falling to 0.481 (0.143–1.000) with 21.1% outliers at 75%, and 0.143 (0.000–0.538) with 53.8% outliers at 50%, despite cosine remaining high (median 0.999950 at 50%). Contamination mixing yields a similar escalation: at 5% mixing, outliers remain near baseline (3.1%) with Jaccard 0.667 (0.600–0.818), but at 10% mixing outliers rise to 15.7% and Jaccard falls to 0.538 (0.379–0.667), and at 20% mixing outliers reach 56.4% with Jaccard 0.290 (0.143–0.481). Species-stratified summaries show that *Listeria* and *B. subtilis* are among the most sensitive under contig dropout and contamination, whereas *Salmonella* and *E. coli* O157:H7 retain comparatively higher neighborhood stability. Together, these curves establish an actionable calibration between assembly quality/contamination and expected neighborhood drift.

**Figure 8.**
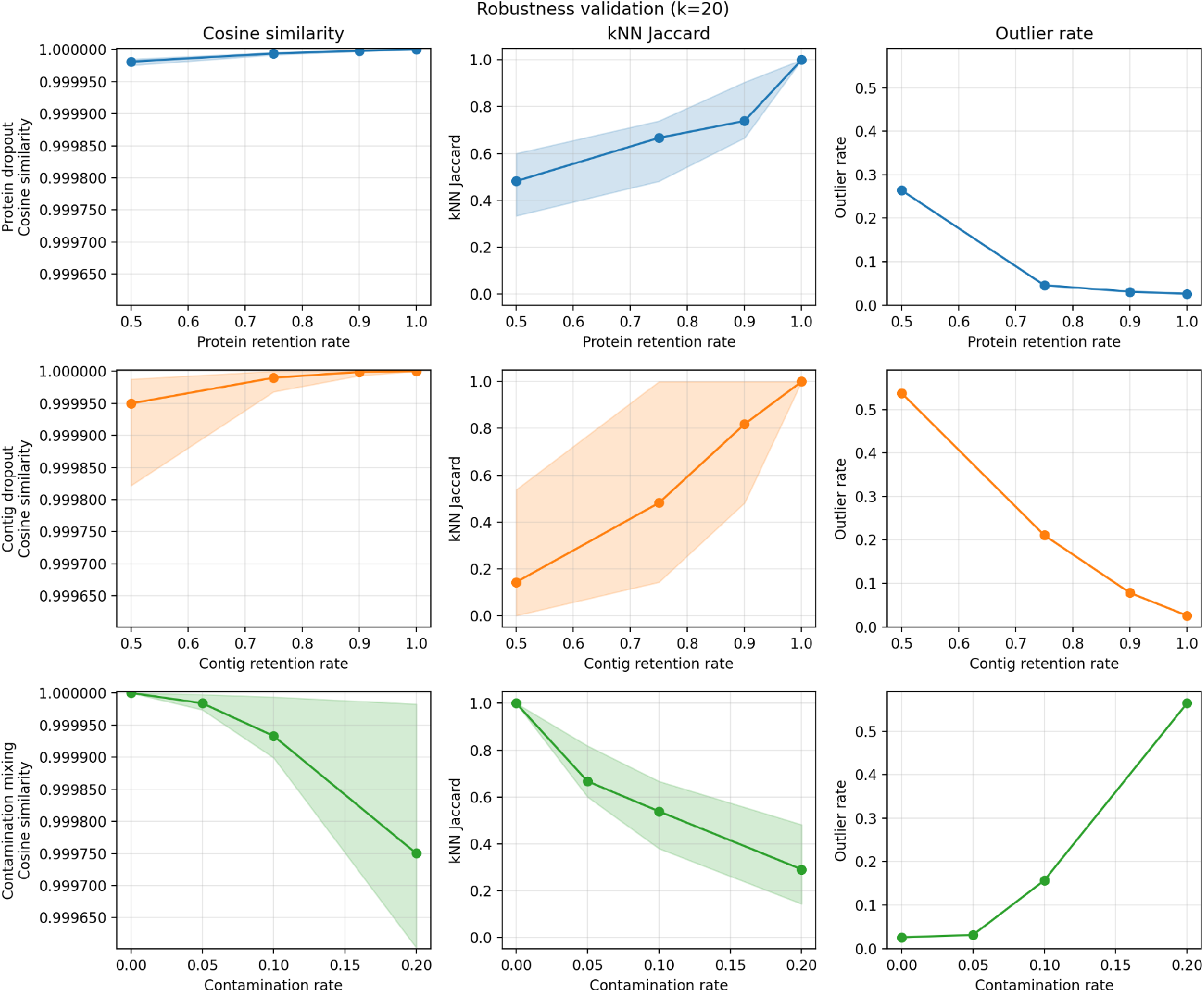
Robustness validation under missingness and contamination. Robustness validation panels (k=20). A balanced sample of 200 genomes per species (n=1,800) is used, with three random repeats per rate (n=5,400 per rate). Rows correspond to protein dropout (top), contig dropout (middle), and contamination mixing (bottom). Columns report cosine similarity to the full embedding (median with IQR band), kNN neighborhood stability (Jaccard overlap), and outlier rate using the reference kNN distance threshold (Q3 + 1.5×IQR). Contamination mixing replaces 5%, 10%, or 20% of proteins with proteins sampled from a different species.

## 4. Discussion

Having established that whole-proteome ESM-2 embeddings recover taxonomic structure, yield high-purity clusters, and provide actionable confidence signals, we now interpret these findings in the context of bio-surveillance requirements, position the approach relative to state-of-the-art methods, and outline the path toward deployment.

### 4.1 What Mean-Pooled Whole-Proteome Embeddings Encode

The strong low-dimensional structure and high species homophily indicate that whole-proteome, mean-pooled ESM-2 genome embeddings capture dominant, taxonomically aligned signals, which is consistent with the view that PLM representations encode broad evolutionary and functional patterns at the protein level [4,5]. For bio-surveillance, this is operationally valuable in the sense that taxonomy-aligned neighborhoods can support rapid retrieval and coarse prioritization without alignment, while geometry-derived diagnostics (homophily, purity, outliers) provide a principled way to expose uncertainty and flag boundary cases for escalation. To be precise, our “context-free” claim applies *at the genome level*: ESM-2 is contextual within each protein sequence, but mean pooling discards gene order, co-localization, and operon-scale context. These results also align with evidence that medium-sized protein language models transfer well on realistic downstream tasks [17].

### 4.2 Positioning Relative to State of the Art

The landscape of computational genomics offers several alternative approaches to genome representation, each with distinct trade-offs. Alignment-free sketching tools such as Mash have become foundational for large-scale genomic indexing, providing very fast nucleotide-level similarity search [3]; containment-based variants like Mash Screen are arguably closer production competitors for rapid detection in mixed samples [23]. Our approach is computationally heavier at embedding time, but produces compact vectors that encode protein-level functional and structural signals beyond raw nucleotide identity [5]. This trade-off becomes favorable in a cache-first setting where embeddings are computed once and reused across downstream analytics and models.

The efficiency profile of our approach merits explicit consideration. At inference time, genome representation requires per-protein embedding plus constant-time averaging, and once cached it enables fast CPU-feasible retrieval, scoring, and re-analysis. Contextual genome language models that explicitly model gene order or gene–gene interactions must operate over thousands of tokens per genome, with attention costs that can scale quadratically in sequence length. This does not diminish the value of contextual models, but rather, it clarifies why a cache-first mean-pooling baseline serves as a useful operational substrate: it amortizes expensive embedding computation and provides geometry-aware diagnostics that can determine when more expensive contextual modeling is warranted.

At the nucleotide level, foundation models such as DNABERT/DNABERT-2 [10,11], Nucleotide Transformer [9], HyenaDNA [12], and GenSLMs [13] can in principle avoid gene calling and capture noncoding signals. However, these models face long-context challenges on megabase-scale bacterial genomes [6] and often focus on specific organismal regimes (human genomics [9,10,11,12], viruses [13]). Protein-level representations reduce sequence length and emphasize functional units, offering a complementary route for microbial surveillance. Emerging long-range architectures partially address these scaling limitations: GENA-LM provides long-sequence DNA foundation models [22], while state-space models with biological inductive biases such as reverse-complement equivariance (e.g., Caduceus) aim for O(n) scaling [21]. These approaches hold promise for future foodborne surveillance, but bacterial whole-genome benchmarks and operational pipelines remain less standardized than in human genomics.

Among protein- and gene-level genome language models, several recent contributions define the current frontier. gLM contextualizes ESM-2 protein embeddings at the operon/contig scale (∼30 genes) to model co-regulation and function [7]. Protein Set Transformer aggregates proteomes as sets to build viral genome language models for viromics [8]. BacPT and Bacformer extend this direction toward whole-proteome or ordered-genome context modeling in bacteria [18,19]. Our mean-pooling baseline is deliberately simpler than these models, yet it serves a practical role: it delivers a scalable cache and artifact pipeline for producing training and analysis data, exposes a geometry-aware diagnostic layer (homophily, purity, outliers) that can operate before or alongside contextual models, and provides a clear baseline for quantifying what additional modeling capacity is required to move beyond taxonomy toward within-species virulence discrimination.

In terms of scale, our dataset (21,657 complete genomes) is substantial for a research-level geometry study and complements larger pretraining corpora used in foundational models (e.g., gLM’s metagenomic contigs [7] and bacteria-scale genome corpora in Bacformer [19]). Looking ahead, newer protein foundation models such as ESM3 [24] suggest that protein-level representations will continue to improve, further strengthening the case for cache-first reuse across bio-surveillance pipelines.

### 4.3 Toward Operational Deployment: A Bio-Surveillance-Oriented Framework

Our long-term goal is to translate these embedding-space diagnostics into an end-to-end system for food safety triage, complementing emerging FAIR and adaptable workflows for foodborne pathogen surveillance [25]: a cache-first pipeline that embeds new assemblies once, then performs fast retrieval, scoring, and evidence-driven escalation under operational constraints. The value of mean-pooled whole-proteome embeddings lies not only in their compact representation, but in the principled, model-agnostic confidence signals their geometry provides.

In practice, homophily and cluster purity serve as deterministic confidence indicators. When a query genome falls in a high-consistency neighborhood, retrieval-based triage can proceed as low-risk for coarse routing (e.g., selecting which reference set to prioritize). When homophily degrades with neighborhood scale (Figure 5), the system can explicitly flag these cases as boundary conditions and route them to deeper analysis, including context-aware models, targeted evidence matching against virulence/AMR gene databases, or expert review, rather than returning a brittle high-confidence label. The within-genus risk–coverage curves (Figure 6) demonstrate how this geometry translates into an operator-facing “defer vs decide” policy with quantifiable trade-offs.

Crucially, because our pathogenicity labels are priors, this escalation logic is fundamentally conservative: it is designed to prevent overinterpretation when embedding neighborhoods become label-mixed, not to claim isolate-level virulence resolution from a species-derived proxy.

Finally, strict outliers and distributional shifts in embedding space provide a complementary system-level signal for quality control and novelty monitoring. In operational deployments, these genomes can be prioritized for assembly/annotation checks, contamination screening, and drift tracking, and can trigger recalibration or retraining when they accumulate over time.

The robustness validation (Figure 8) makes this actionable for the system we are envisioning. By jointly stress-testing protein loss, contig fragmentation, and cross-species mixing, it quantifies when local neighborhoods destabilize and when outlier rates spike, enabling calibrated “trust vs defer” rules tied to assembly quality and contamination risk. Because the analysis is driven by the cached per-protein embeddings, it can be rerun whenever the cache expands or when the pipeline ingests new taxa, providing a living calibration layer for surveillance deployment rather than a one-time benchmark.

### 4.4 Limitations

Several limitations bear on both scientific interpretation and any bio-surveillance-oriented deployment.

First, our pathogenicity labels are species/pathotype-derived priors, not isolate-level phenotypes. Consequently, the strong neighborhood agreement we observe largely reflects taxonomic coherence, and zero-shot transfer of the binary label across held-out taxa is unreliable by construction. The within-genus stress tests should therefore be interpreted as pathotype-level discriminability under known annotations, not as discovery of novel virulence factors in unlabeled species.

Second, mean pooling discards protein order and neighborhood context, precisely the structure that encodes pathogenicity islands, operons, and mobile element signatures. These are exactly the regimes where contextual genome language models are expected to add value (e.g., gLM [7], Bacformer [19]).

Third, the representation inherits sensitivity to upstream annotation and proteome composition. Protein extraction depends on gene calling/annotation, and unequal proteome sizes can bias a mean-pooled summary. Empirically, protein count correlates with embedding “edge-case” signals in this dataset (e.g., Spearman ρ = 0.29 between protein count and k-neighbor distance, and ρ ≈ −0.20 between protein count and label homophily), and strict outliers have larger median protein counts than non-outliers (4,902 vs 4,403). These patterns are consistent with a mix of biological diversity and assembly/annotation artifacts, and motivate explicit QC and missingness stress tests. We address this directly with the robustness validation (Sections 2.8 and 3.9), which quantifies how incomplete or contaminated proteomes perturb neighborhood stability and outlier rates. These stress tests are simulated (random protein/contig removal and cross-species mixing) and therefore cannot replace real-world completeness/contamination measurements; they provide a principled stress benchmark but should ultimately be anchored to external QC tools and independent datasets.

Fourth, our dataset is imbalanced (7,000 *Salmonella* vs 200 *C. freundii*; 35:1), which can bias centroid estimates, neighbor mixing asymmetry, and density-based clustering toward abundant taxa. Finally, our cache configuration uses a maximum protein sequence length (`max_prot_seq_len=1024`), so very long proteins may be truncated; the downstream impact has not been quantified here.

We view these limitations as an honest boundary of what a mean-pooled baseline can claim, and as a roadmap for the next iteration: evidence integration (virulence/AMR gene matching), deeper outlier validation (e.g., completeness/contamination checks), and systematic calibration against longitudinal data before making stronger production claims. More broadly, they motivate context-aware genome language models and evidence-driven pipelines as the next step.

### 4.5 Future Directions

Several follow-on analyses are feasible without additional embedding runs and would directly strengthen the path from descriptive geometry to deployment readiness. One direction is to calibrate homophily and cluster purity into empirical decision thresholds by mapping them to retrieval error rates (or downstream misclassification) under held-out splits. A second is to stress-test cross-taxon generalization by training on a subset of taxa and evaluating on held-out taxa, quantifying how much of the signal is genuinely transferable beyond taxonomy priors. Third, robustness-to-missingness has been implemented via protein and contig dropout plus contamination mixing using the cached per-protein embeddings; extending this to assembly-quality stratification and external validation datasets would further harden the pipeline. Finally, an interactive “hard-case” atlas of low-homophily and strict-outlier genomes, paired with nearest neighbors and metadata, would support human-in-the-loop quality control and create a concrete bridge between embedding diagnostics and actionable bio-surveillance workflows.

## 5. Conclusions

Whole-proteome ESM-2 embeddings provide a scalable, cache-friendly representation that strongly recovers microbial taxonomy and yields the three actionable signals central to our bio-surveillance framework (Figure 1): confidence scores derived from neighborhood homophily, triage reliability via cluster purity, and early warnings through outlier detection. At the same time, moderate agreement with a binary pathogenicity prior underscores the need for context-aware genome language models to advance from taxonomy-aligned retrieval toward strain-level virulence discrimination and evidence attribution [7,18,19].

For operational food safety surveillance, these results establish a concrete, deployment-aligned foundation: a reusable embedding cache paired with geometry-derived confidence and novelty signals that can gate retrieval, prioritize evidence extraction, and escalate boundary cases to richer models or expert review.

## Data and Code Availability

All code, analysis notebooks, and exported artifacts are available upon request from the authors.

